# A Convection-Free Apparatus for Low Force and High Efficiency Centrifugation

**DOI:** 10.1101/2023.08.18.553772

**Authors:** Teng-Chieh Yang

## Abstract

Centrifugation is a common technique for material separation. Most biological materials are separated under a relatively strong centrifugal force. Despite several reports highlighted the impact from centrifugal force on the separated materials, low force centrifugation typically results in low efficiency of separation and may not be achievable from the traditional centrifugation apparatus. Here, we present a novel apparatus for separating materials under low centrifugal force with high efficiency. Compare to the traditional apparatus, the design of this apparatus minimizes the convection during centrifugation. Using homogeneous polystyrene particles, we confirmed that the centrifugation process is essentially convection-free. Separation of human blood cells indicated that, under a fixed centrifugal force, less than half of the centrifugation time is required compared to the traditional apparatus. The isolated human peripheral blood mononuclear cells have higher viability and can afford multiple centrifugations without impacting cell viability. Additionally, this new apparatus has a higher resolution to separate materials by size and is sensitive to detect size heterogeneity. This apparatus provides a new opportunity for efficient centrifugation under low centrifugal force and for a systematic evaluation of the impact from centrifugal force on the quality of separated materials.

## INTRODUCTION

Centrifugation is one of the most common techniques for separating materials of interest in chemical, dairy, agricultural, pharmaceutical, and medical industries^1–5^. The separation is facilitated in a liquid medium where materials are separated by the centrifugal force. In a medium with a homogeneous density, materials with different densities can be separated predominantly by their size and shape (i.e. differential sedimentation)^1^. In a medium with a density gradient, materials can be separated to specific density layers matching their own densities (i.e. isopycnic separation)^1^.

The application of centrifugation can be classified into two categories: analytical and preparative. Analytical ultracentrifugation (AUC) is an analytical tool to study the size, shape, and interactions of molecules under ultra-high centrifugal forces^6–8^. It has sector-shaped sample chambers horizontally fitted in a rotor, allowing molecules to sediment along the direction of centrifugal force until their reach the bottom of the chamber. The design of AUC provides the sensitivity to characterize biological molecules like proteins or nucleic acids, but is not suitable for material preparation. Preparative centrifugation typically has tubes or bottles to accommodate a larger sample volume. These tubes or bottles have parallel walls to be fitted into a fixed angle or swinging bucket rotor. The design of preparative centrifuge tubes and fixed angle rotor drives materials to collide with the wall and each other during centrifugation, creating shear stress on the materials and *convection* in the medium^9, 10^. Convection further reduces the efficiency of separation, thereby forcing the materials to be separated under a stronger centrifugal force or for a longer centrifugation time.

Typical preparative centrifugation for biological materials (e.g. tissues, cells, organelles, proteins, or nucleic acids) has been conducted under centrifugal force ≥ 100 *g*, equivalent to having materials subjected to ≥ 1,000 meter under the water. Not surprisingly, it has been reported that the centrifugal force has measurable impacts on the separated materials^10–14^. For example, increased hemolysis coupled with ATP release from red blood cells (RBC) has been correlated with centrifugal force and centrifugation time^12^. Additionally, the integrity of adipose tissue was damaged under strong centrifugal force, which was attributed to a lower success rate of fat grafting compared to the non-centrifugation method^14^. Furthermore, centrifugal force has been shown to alter the surface properties of bacteria together with their interior structures^13^.

Despite these reports, few studies systematically evaluated the impact of centrifugation on the quality of separated materials, likely due to the lack of a centrifugation apparatus that can separate materials under low centrifugal force. Here, a new preparative apparatus was designed for high efficiency separation under the low centrifugal force (6 *g* - 25 *g*). The key feature of the design is to minimize convection during centrifugation, similar to the design of AUC but under low force. The performance of this apparatus was tested with highly homogenous polystyrene particles (PSP) and confirmed that these particles were sedimented in the convection-free (CF) environment. Compared to the traditional centrifugation, this apparatus reduces the time of separation by more than 50% under the same centrifugal force and provides the sensitivity to detect sample heterogeneity. Importantly, this apparatus substantially improves the quality of separated materials such as cell viability even under repetitive centrifugations. Therefore, this newly designed apparatus offers an opportunity for systematic evaluation of low force centrifugation and has the potential to improve the quality and consistency of materials prepared in the research and clinical laboratories.

## METHODS

### General Reagent Details

Five (5) μm white polystyrene particles (PSP) were from ThermoScientific (Duke standards). One (1) μm yellow PSP and 6 μm red PSP were from Polyscience Inc. Dulbecco’s phosphate-buffered saline (DPBS) was from Sigma-Aldrich. Polysorbate 20 (PS-20) was from J.T. Baker. Trypan blue staining solution was from Corning. Ficoll-Paque was from GE healthcare.

### General Experimental Details

All centrifugation experiments were conducted in an Eppendorf 5702R centrifuge with a swing bucket rotor (Eppendorf, A-4-38) at 20°C. All samples were centrifuged with slow acceleration and no break during deceleration.

### Model Design and 3D-Printing for Convection-Free (CF) Apparatus

The CF tubes and adaptors were designed using Inventor from Autodesk. The adaptors were 3D-printed layer-by-layer with 0.1 mm per layer under UV cured condition using a RFS600 Pro printer from RealFun. After 3D-printing, the adaptors were washed with isopropyl alcohol followed by polishing. The CF tubes were 3D-printed layer-by-layer with 0.05 mm per layer under UV cured condition using a Form 3 3D-printer from Formlabs. After 3D-printing, the tubes were washed with isopropyl alcohol followed by coating and polishing.

### Characterization of Polystyrene Particles (PSP) Sedimentation

The sedimentation process of polystyrene particles (PSP) was qualitatively monitored by visual analysis of the centrifuge tubes followed by quantitative concentration measurements. After each centrifugation time point, 50 uL of the sample was taken from the top and the middle of solution and then mixed for the absorbance measurement at wavelength 405 nm using SpectraMax (Molecular Devices). Absorbance at this wavelength is proportional to the concentration of white PSP (from 0.015 % to 0.15 %, w/v). Additionally, yellow and red PSP have peak absorbance at wavelengths 415 nm and 550 nm, respectively. Thus, in the mixture of yellow and red PSP, the concentration of each PSP can be determined from the dual-wavelength approach (DWA) described in the following section. All PSP sedimentation experiments were performed in 1X DPBS with 0.01% (w/v) PS-20.

### PSP Concentration Determination by Dual-Wavelength Approach (DWA)

In the yellow and red PSP mixture, the absorbance at wavelengths 415 nm (*A*_415_) and 550 nm (*A*_550_) is the sum of absorbance from each PSP (*A*_*y*,415_, *A*_*y*,550_ for yellow PSP and *A*_*r*,415_ *A*_*r*,550_ for red PSP) and can be expressed as:

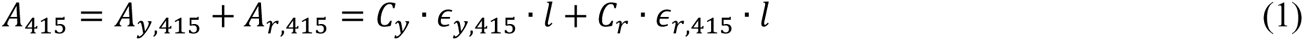

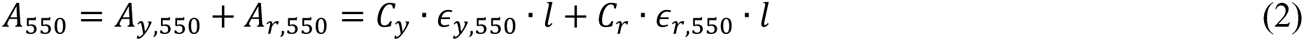

where *C*_*y*_ and *C*_*r*_ are the concentrations of yellow and red PSP, respectively; *є*_*y*,415_ and *є*_*r*,415_ are the extinction coefficients of yellow and red PSP at wavelength 415 nm, respectively; *є*_*y*,550_ and *є*_*r*,550_ are the extinction coefficients of yellow and red PSP at wavelength 550 nm, respectively, and *l* is the path length.

Rearranging Equation (1) and (2), *C*_*r*_ and *C*_*y*_ can be expressed as a function of the total absorbance and the extinction coefficients for two PSP at two wavelengths :

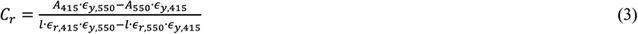

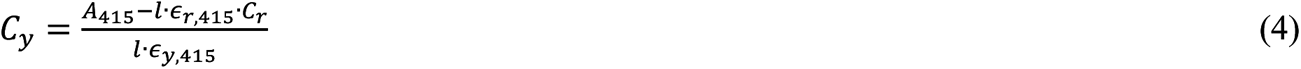

where the extinction coefficients were determined from the known concentrations of red and yellow PSP: *є*_*y*,415_=137.7 cm^-1^ % (w/v)^-1^; *є*_*y*,550_=122.9 cm^-1^ % (w/v)^-1^; *є*_*r*,415_=11.5 cm^-1^ % (w/v)^-1^; *є*_*r*,550_=12.3 cm^-1^ % (w/v)^-1^. Therefore, *C*_*r*_ and *C*_*y*_ can be calculated from the total absorbance measurements at wavelengths 415 nm and 550 nm.

To confirm the accuracy of DWA, varying concentrations of red PSP (0.08%, 0.04%, and 0.02%, w/v) were mixed with 0.01% (w/v) yellow PSP followed by the total absorbance measurements. The concentrations of individual PSP were determined from Equation (3) and (4) using the above extinction coefficients. All PSP concentrations determined by DWA were within 15% of the known concentrations.

### Preparation of Human Peripheral Blood Mononuclear Cells (PBMC)

Human blood sample was obtained from the author itself and subsequently used for PBMC isolation following the procedure described elsewhere^15^. All methods were carried out in accordance with relevant guidelines and regulations. Briefly, equal volume of anti-coagulated blood and 1X DPBS were mixed and then layered on top of Ficoll-Paque in a centrifuge tube. The mixture was centrifuged at 400 *g* for 40 min. The isolated PBMC were transferred to a new tube, washed twice with 1X DPBS and then resuspended in 30 mL 1X DPBS to a final concentration of approximately 5E+5 cell/mL determined from the hemocytometer. The viability of PBMC was determined from Trypan blue staining as described elsewhere^16^. The statistical difference of cell viability was evaluated by the paired t-test.

## RESULTS AND DISCUSSION

### The Design of Convection-Free (CF) Apparatus

The CF apparatus is composed of a CF tube and an adaptor. To minimize convection during centrifugation, the CF tube was designed with the walls aligned with the centrifugal force and a flat bottom for materials to settle without generating turbulence. This flat bottom also enables sedimented materials to be gently resuspended without the need of pipetting which is known to generate shear stress (Figure 1)^17^. To observe the sedimentation process, the CF tube was 3D-printed with clear acrylonitrile butadiene styrene - like resin followed by coating and polishing (see Methods).

**Figure 1.**
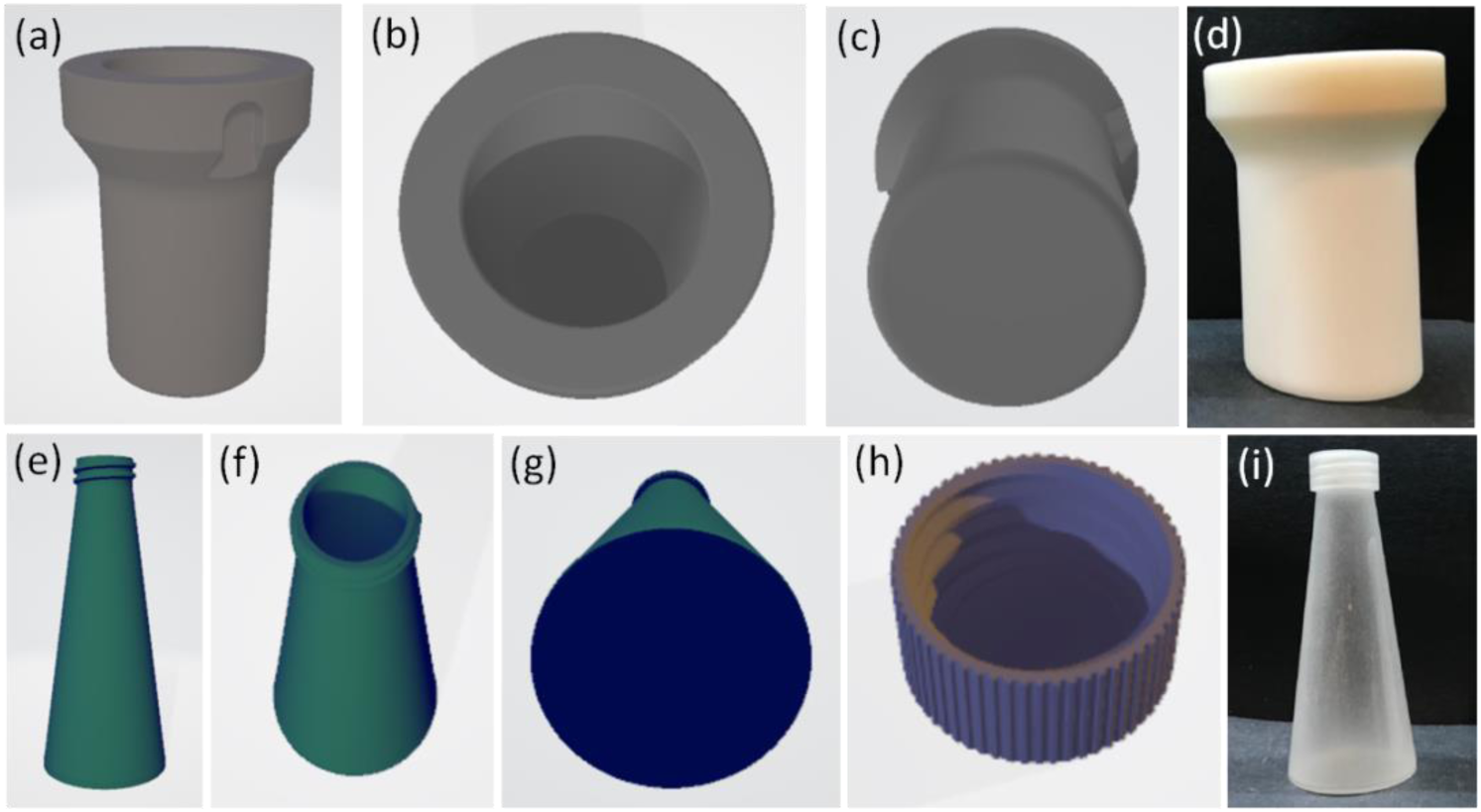
Design of CF Apparatus. (**a**) - (**c**) Design of CF tube adaptor. (**d**) 3-D printed CF tube adaptor. (**e**) - (**h**) Design of CF tube. (**i**) 3-D printed CF tube.

Traditional aluminum swinging bucket for hosting the centrifuge tube is heavy and inefficient for low force centrifugation. A light-weighted adaptor was designed for directly holding a CF tube during centrifugation. This adaptor was 3D-printed with polymer resin and is 30% lighter than the traditional aluminum bucket, allowing a near horizontal swinging plan under low centrifugal forces (≤ 10 *g*).

### Proof of Concept with Polystyrene Particles (PSP)

To test the design of CF apparatus, highly homogenous and spherical PSP with a diameter of 5 µm were sedimented in the following condition: 20 mL of 0.1% (w/v) PSP in 1X DPBS with 0.01% (w/v) PS-20 under 25 *g.* With 20 mL in a CF tube, the solution height is approximately 2.3 cm - the longest distance PSP have to travel before reaching the bottom. Given the sedimentation velocity of 5 µm PSP under 25 *g* is approximately 0.077 cm/min (Table 1 and Supplemental), the estimated time to sediment all 5 µm PSP in the CF environment is 29.9 min (29.9=2.3/0.077).

**Table 1.** Physical and Hydrodynamic Properties of Polystyrene Particles (PSP) The apparent sedimentation coefficient (*s\**) is calculated from the sedimentation in 1X DPBS and 0.01% (w/v) PS-20 with the solution density of 1.005 g/mL and viscosity of 0.01 g cm^-1^ sec ^-1^ at 20°C and using partial specific volume of 0.956 mL/g for polystyrene^20^.

| Molecules | Radius (cm) | Density (g/mL) | $s^*$ (S) | Sedimentation velocity under 6 g (cm/min) | Estimated time to sediment 20 mL in CF environment under 6 g (min) | Sedimentation velocity under 25 g (cm/min) | Estimated time to sediment 20 mL in CF environment under 25 g (min) |
| --- | --- | --- | --- | --- | --- | --- | --- |
| 1 μm PSP | 5.00E-5 | 1.05 | 2.27E+4 | 7.7E-4 | 2990.1 | 3.1E-3 | 747.5 |
| 5 μm PSP | 2.50E-4 | 1.05 | 5.69E+5 | 0.019 | 119.6 | 0.077 | 29.9 |
| 6 μm PSP | 3.00E-4 | 1.05 | 8.19E+5 | 0.028 | 83.1 | 0.111 | 20.8 |

To monitor the sedimentation process of 5 μm PSP under 25 *g*, visual analysis was performed after centrifugation for 10, 20, 40, and 60 min. After 20 min of centrifugation, solution in the CF tube appeared to be more translucent compared to the traditional Falcon^TM^ tube (Figure 2). Even after 60 min of centrifugation, solution was still turbid in the Falcon^TM^ tube, indicating PSP remained in suspension. The concentration measurement of PSP also suggested that, after 20 min of centrifugation, only 7.6% PSP remained in solution from the CF tube compared to 29.8% in solution from the Falcon^TM^ tube. Additionally, regression analysis was applied to determine the kinetics of PSP sedimentation and indicated that less than 3% PSP remained in solution from the CF tube after 30 min of centrifugation (Figure 2b). This is consistent with the sedimentation time (29.9 min) estimated from the sedimentation velocity, suggesting that centrifugation in this apparatus is essentially convection-free.

**Figure 2.**
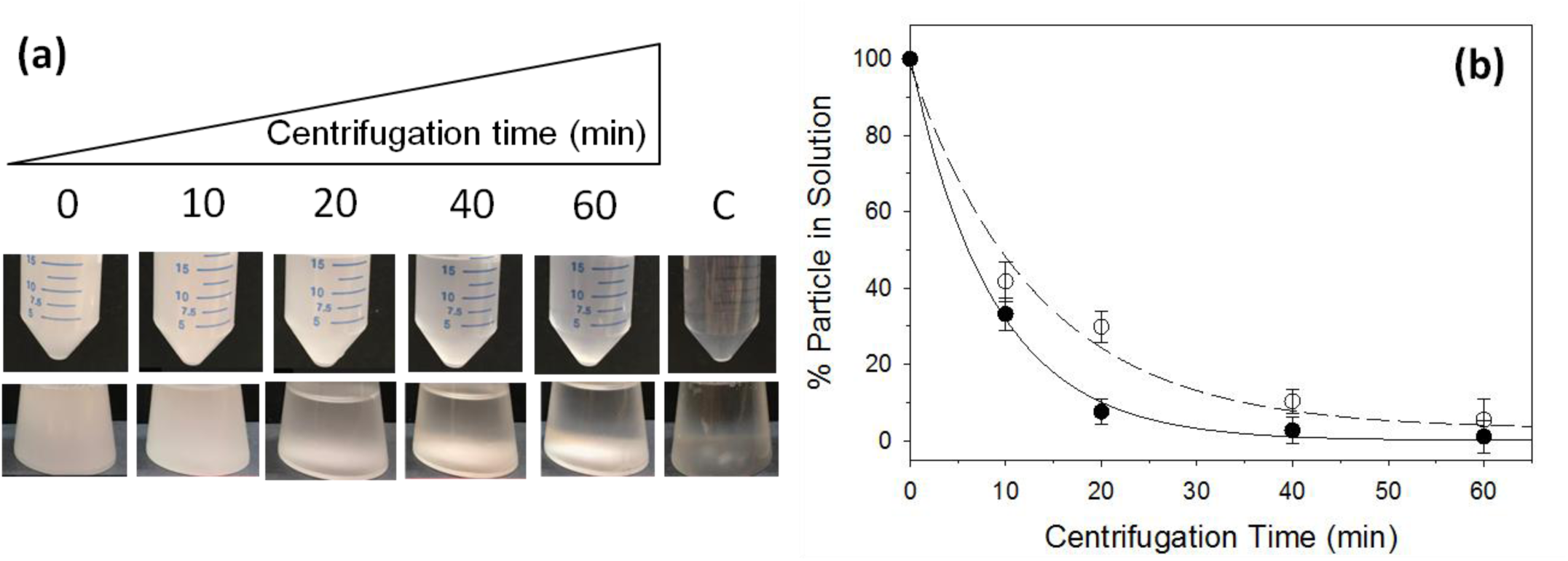
Sedimentation of 5 µm polystyrene particles (PSP) in the CF Apparatus. (**a**) Visual analysis of 5 µm PSP (0.1%, w/v) sedimentation under 25 *g* for up to 60 min. C is the control without PSP. (**b**) Quantitative analysis of percent 5 µm PSP in solution after sedimentation. Black circles: CF tube. White circles: Falcon^TM^ tube. Solid and dotted lines are the non-linear least square (NLLS) analysis using the following equation: Y=A*exp(-k*t)+B where Y is the percent particle in solution, A is the amplitude, k is the kinetic constant, B is the baseline, and t is the centrifugation time. During the NLLS analysis, A, B, and k were allowed to float until finding the best fit parameters. Error bars indicate the standard deviation from triplicates.

### CF Apparatus Enables Low Force Separation

The typical centrifugal force for separating eukaryotic cells is 250 – 800 *g*, prokaryotic cells is 1,000 – 8,000 *g,* organelles is 20,000 – 150,000 *g*, and proteins/oligonucleotides is 60,000 – 200,000 *g*^18^. These centrifugal forces are higher than the minimum required forces for separation due to the low separation efficiency inherent in the design of traditional apparatus.

To determine the feasibility of CF apparatus under ultra-low centrifugal force, 20 mL of 5 μm PSP was sedimented under 6 *g*. After 120 min centrifugation, more than 98% PSP were sedimented in solution from the CF tube (Figure 3). This is consistent with the sedimentation time estimated from the sedimentation velocity under 6 *g* - 121 min (121=2.3/0.019). In contrast, more than 10% particles remained in solution after 120 min centrifugation in the Falcon^TM^ tube, suggesting that the design of traditional tubes is inefficient to separate materials under this ultra-low centrifugal force.

**Figure 3.**
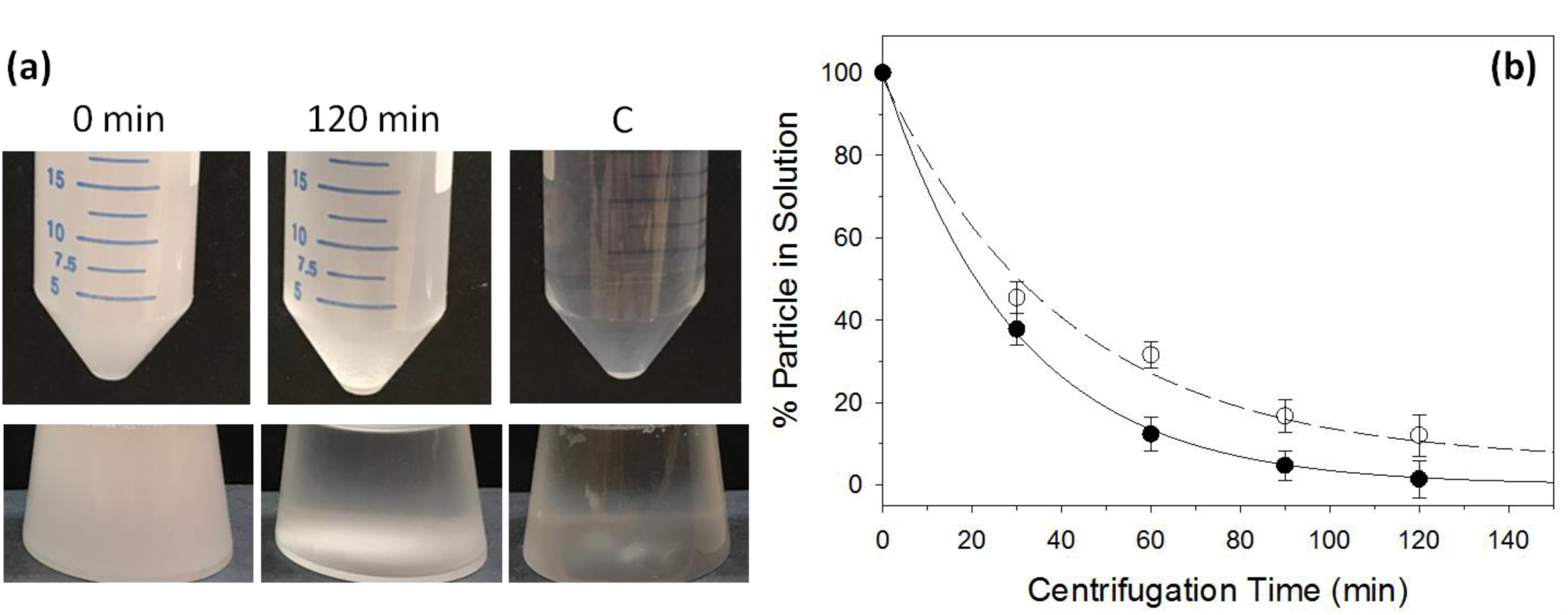
Material separation under low centrifugal force. (**a**) Visual analysis of 5 µm PSP (0.1%, w/v) sedimentation under 6 *g* for up to 120 min. C is the control without PSP. (**b**) Quantitative analysis of percent 5 µm PSP in solution after sedimentation. Black circles: CF tube. White circles: Falcon^TM^ tube. Solid and dotted lines are the NLLS analysis using the following equation: Y=A*exp(-k*t)+B where Y is the percent particle in solution, A is the amplitude, k is the kinetic constant, B is the baseline, and t is the centrifugation time. During the NLLS analysis, A, B, and k were allowed to float until finding the best fit parameters. Error bars indicate the standard deviation from triplicates.

### High Efficiency Separation of Biological Samples

The sedimentation of human peripheral blood mononuclear cells (PBMC) was evaluated in the CF and Falcon^TM^ tubes. The experiments were conducted under three centrifugal forces, 6 *g*, 25 *g*, and 157 *g* with a fixed centrifugation time - 15 min. Under higher centrifugal forces typically used for PBMC preparation (≥ 157 *g*), no difference on the total PBMC recovery was observed. However, under lower centrifugal forces (e.g. ≤ 25 *g*), PBMC were recovered with higher efficiency in the CF tube (Figure 4a). Additionally, the viability of PBMC is significantly (p<0.05) improved under low centrifugal forces as discussed in the latter section.

**Figure 4:**
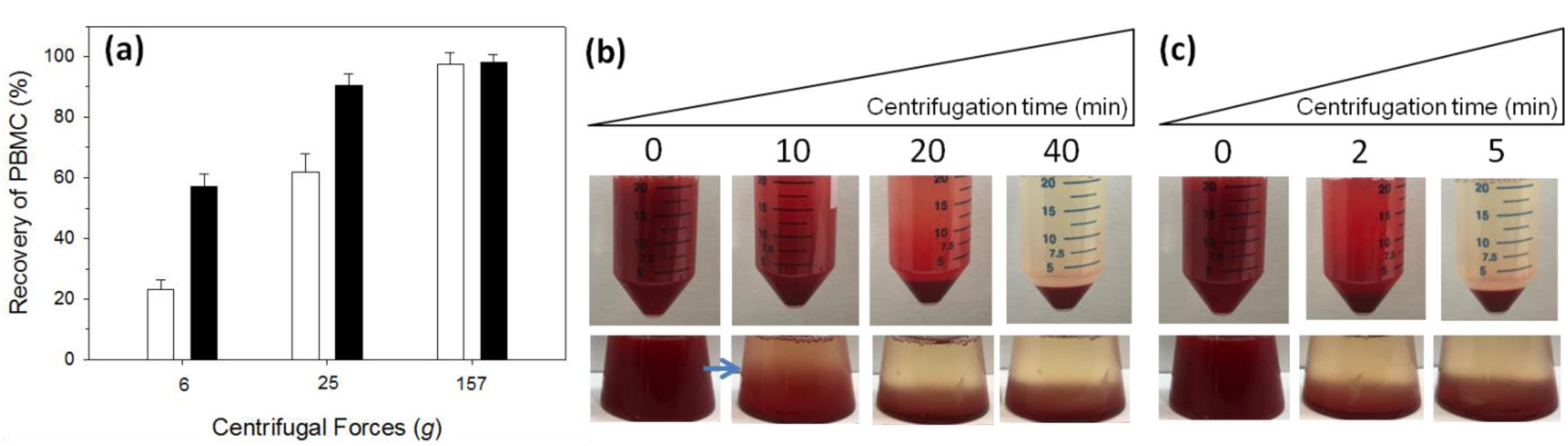
Separation of biological materials. (**a**) Recovery of human PBMC under different centrifugal forces. The recovery is calculated from the total PBMC using a hemocytometer. Black bars: CF tube. White bars: Falcon^TM^ tube. Error bars indicate the standard deviation from triplicates. (**b**) Separation of human plasma under 6 *g*. (**c**) Separation of human plasma under 25 *g*.

In addition to PBMC separation, the isolation of plasma was compared in the CF and Falcon^TM^ tubes under low centrifugal forces, 6 *g* and 25 *g* (Figure 4b and 4c). Visual analysis indicated that, under the identical centrifugal force, plasma was isolated in the CF tube with ≤ 50% of the time required in the Falcon^TM^ tube. Interestingly, the average erythrocyte sedimentation rate (ESR) from adult hematology test is approximately 0.033 cm/min under 1 *g*^19^, translating to 0.20 and 0.81 cm/min under 6 *g* and 25 *g*, respectively (See Supplemental). These rates suggest that 20 mL red blood cells (RBC) should complete sedimentation in 11.5 min and 2.8 min under 6 *g* and 25 *g*, respectively, consistent with the visual analysis in Figure 4b and 4c.

Note, a clear concentration boundary of RBC was observed in the CF tube after 10 min centrifugation under 6 *g* (blue arrow in Figure 4b), suggesting a minimized solution disturbance as compared to a broad concentration gradient of RBC observed in the Falcon^TM^ tube. This is similar to the concentration boundary observed during the sedimentation velocity AUC experiment^6–8^. As such, the design of CF apparatus should provide a higher resolution for material separation by their size.

### Material Separation by Size

A mixture of 6 μm red PSP (0.04%, w/v) and 1 μm yellow PSP (0.005%, w/v) was centrifuged under 25 *g.* Visual analysis indicated that red PSP were separated from yellow PSP after 40 min centrifugation in the CF tube (Figure 5a). During centrifugation, the concentrations of red and yellow PSP were further determined by the dual-wavelength approach (DWA) as described in the Methods. Indeed, after 40 min centrifugation, more than 90% of the red PSP were separated from the yellow PSP in the CF tube, compared to only 76% separation in the Falcon^TM^ tube. Further increased centrifugation time to 60 min increased the separation of red PSP; however, a small portion (5-10%) of yellow PSP also began to sediment under the extended centrifugation time (Figure 5b). To optimize the separation condition, a short-term centrifugation (10 min) under different centrifugal forces was evaluated in the CF and Falcon^TM^ tubes. It was determined that, centrifugation with the CF apparatus under 100 *g*, red PSP were most efficiently sedimented while ≥ 95% of yellow PSP remained in solution (Figure 5c).

**Figure 5:**
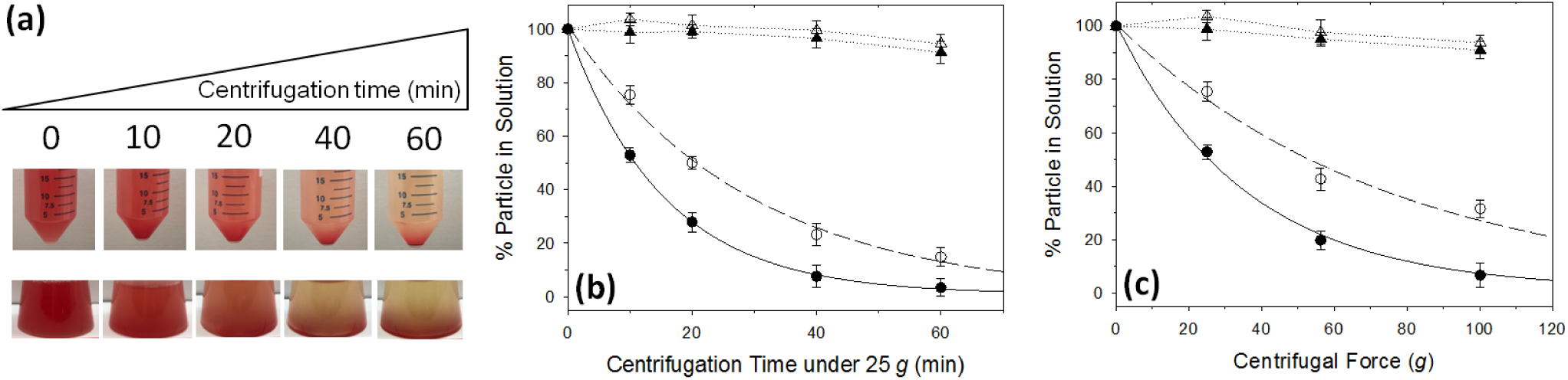
Separation of a PSP mixture. (**a**) Visual analysis of a mixture of red 6 µm PSP (0.04%, w/v) and yellow 1 µm PSP (0.005%, w/v) sedimentation under 25 *g*. (**b**) Quantitative analysis of red and yellow PSP sedimentation under 25 *g* for up to 60 min. (**c**) Quantitative analysis of red and yellow PSP sedimentation for 10 min under different centrifugal forces. Solid and dotted lines are the NLLS analysis using the following equation: Y=A*exp(-k*t)+B where Y is the percent red PSP in solution, A is the amplitude, k is the kinetic constant, B is the baseline, and t is the centrifugation time. During the NLLS analysis, A, B, and k were allowed to float until finding the best fit parameters. Error bars indicate the standard deviation from triplicates.

### Detection of Size Heterogeneity

During the previous separation study, the sedimentation time for 6 μm red PSP was found to be longer than expected (Table 1). Based on the sedimentation velocity of 6 μm PSP under 25 *g* (0.111 cm/min, Table 1), the time to sediment 20 mL is approximately 21 min (21=2.3/0.111). However, it took almost double the time to sediment in Figure 5b and thus can be attributed to the following three reasons: an interaction between 6 μm and 1 μm PSP or self-interaction of 6 μm PSP or the size heterogeneity of 6 μm PSP. The experiment was repeated with 6 μm PSP alone at different concentrations under 25 *g* where longer than expected time of sedimentation was still observed (Figure 6b), indicating the possibility of size heterogeneity. Next, imaging analysis was applied to evaluate the size distribution of 6 μm PSP under light microscope, returning an average size of 6.1 ± 2.3 μm. This size variation (CV=37.7) is larger than the variation of 5 μm PSP (5.0± 0.04 μm, CV=1.0). Therefore, the longer than expected sedimentation time of 6 μm red PSP is a consequence of size heterogeneity. This result also indicated that the CF apparatus is sensitive to detect size heterogeneity and may be useful for determining the sedimentation coefficient of large molecules under low centrifugal forces, which is not achievable by AUC.

**Figure 6:**
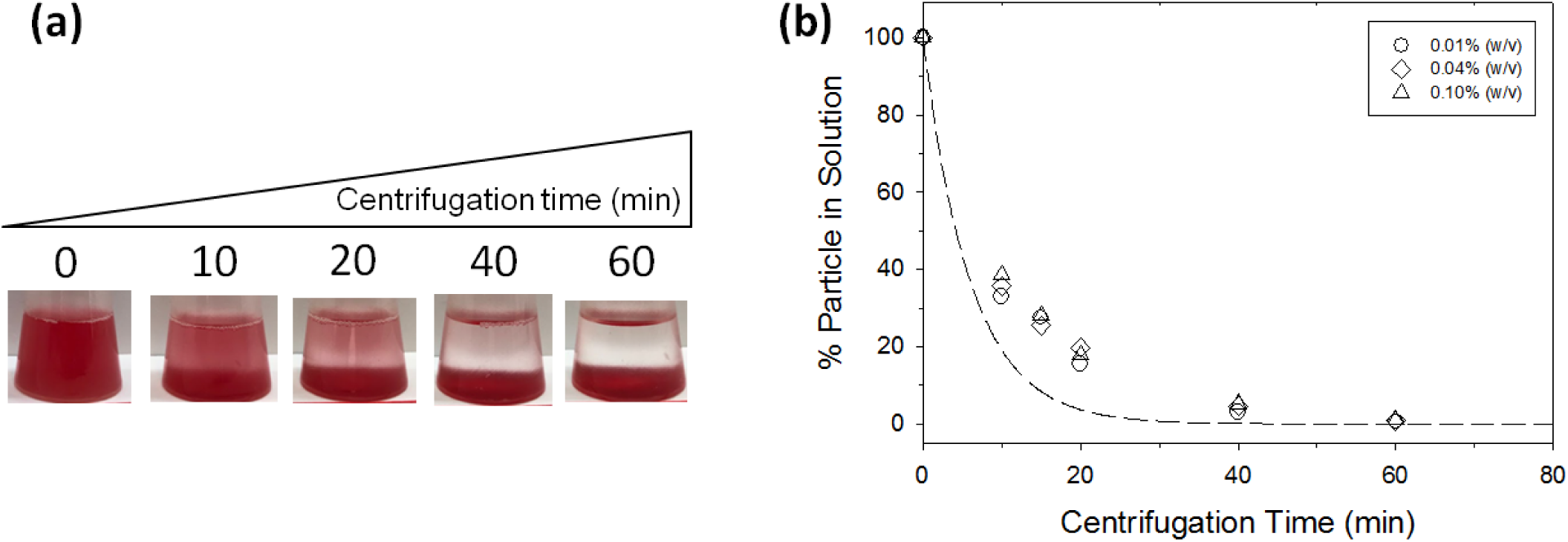
CF apparatus detects size heterogeneity of PSP. (**a**) Visual analysis of 6 µm PSP (0.04%, w/v) sedimentation under 25 *g* for up to 60 min. (**b**) Quantitative analysis of red PSP (0.01 to 0.1%, w/v) sedimentation under 25 *g* for up to 60 min. Dotted curve is the theoretical sedimentation of homogenous 6 µm PSP calculated using the following equation: Y=A*exp(- k*t)+B where Y is the percent red PSP in solution, A is the amplitude, k is the kinetic constant, B is the baseline, and t is the centrifugation time with the following fixed parameters (A=100, B=0, and k=0.165).

### Improved Cell Viability Using CF Apparatus

It has been reported that the centrifugal force created negative impacts on cells such as premature activation of cell surface receptor or the decreased cell viability^6, 13^. Unfortunately, these studies were performed in the traditional apparatus where higher than required centrifugal forces still create shear stress in the control experiments. Here, the viability of human PBMC was tested under low centrifugal forces and with repeated centrifugations. When PBMC were sedimented in the CF tube with centrifugal forces between 6 *g* and 307 *g* for 15 min, no apparent impact on the cell viability was observed as compared to the decreased cell viability in the Falcon^TM^ tubes under higher centrifugal forces (Figure 7a). Similarly, when PBMC were centrifuged under 307 *g* for 15 min and repeated for up to 3 times, no apparent change on the cell viability was observed using the CF tube as compared to the progressive decrease in cell viability using the Falcon^TM^ tube (Figure 7b). This can be explained by at least two reasons: the reduced shear stress in the CF environment and the flat bottom of CF tubes allowing PBMC to settle during centrifugation and to be reconstituted via gentle swirling during re-suspension. This result also suggested that, even under a lower centrifugal force, the design feature of the apparatus can impact the quality of separated materials.

**Figure 7:**
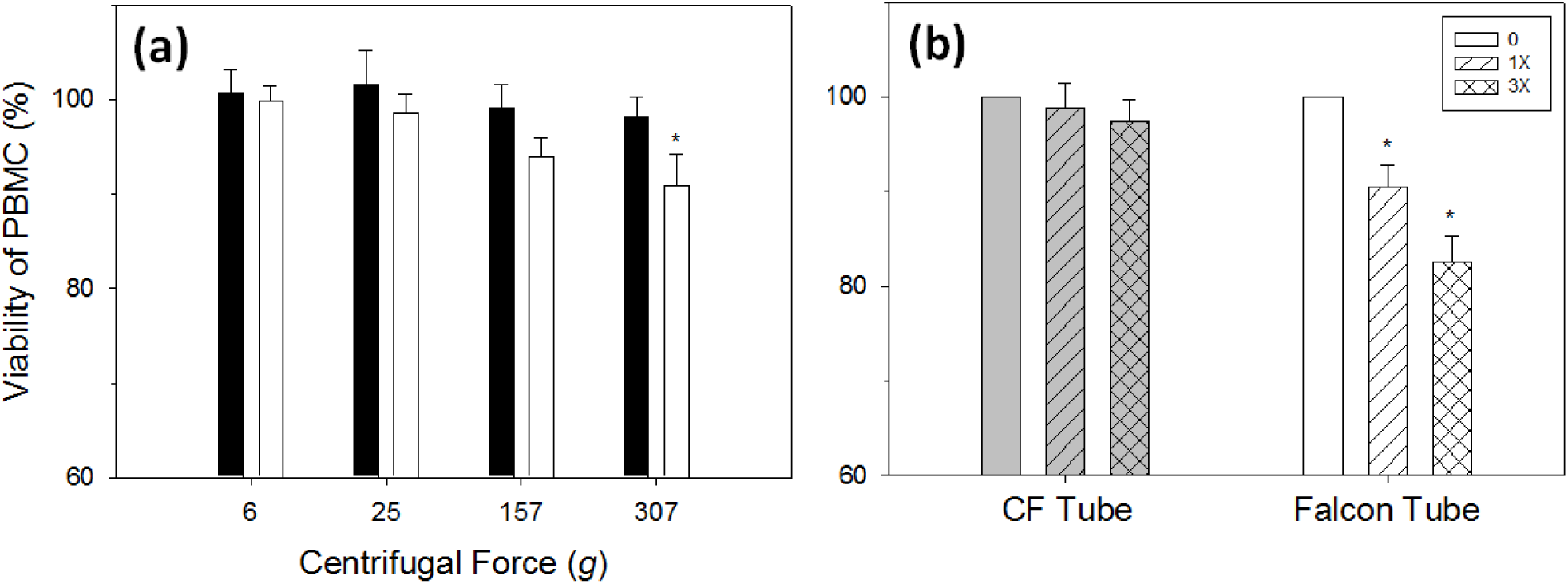
Higher human PBMC viability when separated under CF environment. (**a**) Viability of PBMC under different centrifugal forces for 15 min. Black bars: CF tube. White bars: Falcon^TM^ tube. Asterisks indicates the *p* value of 0.03 calculated from the paired *t-*test between the CF tube and Falcon^TM^ tube under 307 *g* with the *n* of 3 for each condition. (**b**) Viability of PBMC under repeated centrifugations. Each centrifugation is under 307 *g* for 15 min. Error bars indicate the standard deviation from triplicates. Asterisks indicates the *p* value of 0.02 and 0.008 calculated from the paired *t-*test between 0 and 1X, and 0 and 3X, respectively, in the Falcon^TM^ tube with the *n* of 3 for each condition.

## CONCLUSIONS

A novel centrifugation apparatus for low force and high efficiency separation was developed. This is achieved by the design to minimize solution convection during centrifugation. The test with highly homogenous PSP indicated that the time required to sediment is consistent with the time calculated from sedimentation velocity in the CF environment. The material used to construct this apparatus also allows high efficiency separation under low centrifugal force.

When separating human plasma in this apparatus, it took less than half of the centrifugation time required in the traditional apparatus. Additionally, the concentration boundary of RBC was observed during centrifugation, indicating that this apparatus is sensitive to separate materials by size or to detect size heterogeneity for materials with known sizes. This idea was confirmed by separating a mixture of 1 µm yellow and 6 µm red PSP, where an optimum condition - a higher centrifugal force with a shorter duration of centrifugation was identified to efficiently separate 6 µm red PSP from 1 µm yellow PSP. Furthermore, the size heterogeneity of 6 µm red PSP was discovered from longer than expected sedimentation time and confirmed by imaging analysis.

Finally, the design of this apparatus reduces the shear stress induced by the traditional apparatus and thus significantly (*p*<0.05) improves the quality (e.g. viability) of sedimented human PBMC. Centrifugation induced damage on separated materials has been reported but not systematically evaluated. This new apparatus offers an opportunity for evaluating the impact from strong centrifugal force with associated shear stress on the separated materials and for improving the quality and consistency of materials generated from research and clinical laboratories.

## ACKNOWLEDGEMENT

The author would like to thank Wei-Ting Chang for her tireless support and encouragement on this project.

## AUTHOR CONTRIBUTION

T-C.Y. contributed to the entire work including study design, execution, and manuscript preparation.

## DATA AVAILABILITY

The data and models used in this work can be generated from the equations in the Methods section. Contact the corresponding author for the data and materials used in this study.

## COMPETING INTERESTS

The author declares no competing interests.

## Notes

### Competing Interest Statement

The authors have declared no competing interest.

