## Supplemental for "A Convection-Free Apparatus for Low Force and High Efficiency Centrifugation"

Authors: Teng-Chieh Yang*

Affiliations: Teng-Chieh Yang

Corresponding author*

### Supplemental Data

*Calculation of Sedimentation Coefficient and Velocity for Polystyrene Particles (PSP)*

The sedimentation velocity ($v$) of PSP can be expressed as follows:

$$v = s \cdot w^2 r \quad \text{(S1)}$$

where $s$ is the sedimentation coefficient (S); $w$ is the angular velocity (rad/sec), and $r$ is the radial position ($m$). The $s$ of PSP can be expressed as a function of PSP mass ($m$), friction coefficient ($f$), partial specific volume ($\bar{v}$), and solvent density ($\rho$)[1-3]:

$$s = \frac{m(1-\bar{v}\rho)}{f} \quad \text{(S2)}$$

Base on the spherical PSP, $f$ can be expressed as a function of PSP radius ($r_0$) and solution viscosity ($\eta$), and thus Equation (S2) can be further rearranged into:

$$s = \frac{m(1-\bar{v}\rho)}{6\pi\eta r_0} = \frac{2r_0^2 D(1-\bar{v}\rho)}{9\eta} \quad \text{(S3)}$$

where $D$ is the density of PSP. For the 5 μm PSP used in this study, $r_0$ is 2.5E-4 cm, $D$ is 1.05 g/mL, $\bar{v}$ is 0.956 mL/g. With the density and viscosity of 1X PBS and 0.01% (w/v) PS-20 at 20$^{o}$C to be 1.01 g/mL and 0.01 g $\cdot$cm$^{-1}$ $\cdot$sec$^{-1}$, respectively, the apparent $s$ for 5 μm PSP is calculated to be 5.69E+5 S. Centrifugation of PSP under 25 $g$ ($w$=41.89 rad/rev) and with an average radial position from the meniscus and the bottom of solution to be 13.85 cm, $v$ is calculated, using Equation (S1), to be 0.077 cm/min. Therefore, the estimated time to sediment 5 μm PSP in a 20 mL solution with 2.3 cm solution height is approximately 29.9 min (29.9=2.3/0.077).

*Calculation of Sedimentation Coefficient and Velocity for Red Blood Cells (RBC)*

The *s* of RBC can be estimated from Equation (S2) where $f$ is the friction coefficient of RBC. Since RBC has a disk-shape, similar to a oblate spheroid, with approximately 8.1E-4 cm in diameter and 2.0E-4 cm in thickness[4], the axial ratio of 4.1 (4.1=8.1/2.0) translates to a friction coefficient ratio ($f/f_0$) of 1.18, where $f_0$ is the friction coefficient of RBC under the assumption of spherical-shape and can be expressed as a function of the volume ($V$) of RBC and the solution viscosity ($\eta$):

$$f_0 = 6\pi\eta r_0 = 6\pi\eta\left(\frac{3V}{4\pi}\right)^{1/3} \qquad \text{(S4)}$$

Rearranging Equation (S2) and (S4) with $f/f_0$=1.18, the *s* of RBC can be determined from Equation (S5):

$$s = \frac{m(1-\bar{\upsilon}\rho)}{1.18 \cdot 6\pi\eta\left(\frac{3V}{4\pi}\right)^{1/3}} \qquad \text{(S5)}$$

Using Equation (S5) with the mass of RBC (*m*) calculated from its volume (9E-11 mL), density (1.11 g/mL) and $\bar{\upsilon}$ (0.7 mL/g)[5,6], the apparent *s* for RBC in solution is determined to be 4.8E+6 S.

On the other hand, the apparent *s* of RBC can be determined from the average erythrocyte sedimentation rate (ESR): 15 to 30 mm/hr, observed from normal adult samples[7]. This range of ESR is translated to the sedimentation velocity of RBC ($v$): 4.2E-6 to 8.3E-6 m/sec. For acceleration under gravity, Equation (S1) can be rearranged as:

$$s = \frac{v}{a} \qquad \text{(S6)}$$

where $a$ is 9.81 m/sec.

Thus, based on the range of $v$ from ESR, the range of apparent $s$ determined from Equation (S6) is 4.3E+6 S to 8.5E+6 S (i.e. average $s$ is 6.0E+6 S). This is close to the apparent $s$ of RBC calculated from Equation (S5).

Importantly, centrifugation of RBC under 25 $g$ ($w$=41.9 rad/rev) and with an average apparent $s$ to be 6.0E+6 S and an average radial position from the meniscus and the bottom of solution (13.85 cm) in the CF tube, $v$ is calculated to be 1.35E-4 m/sec (i.e. 0.81 cm/min). This is close to the estimated $v$ from the average ESR, 1.39E-4 m/sec, indicating the RBC sedimentation behavior did not change under 25 $g$. At this sedimentation velocity, the expected time to sediment a 20 mL RBC solution is 2.8 min (2.8=2.3/0.81), consistent with the visual analysis in Figure 4C. Similarly, $v$ for RBC is calculated to be 0.20 cm/min under 6 $g$ ($w$=20.9 rad/rev), translating to 11.3 min (11.3=2.3/0.20) for the estimated time to sediment a 20 mL RBC solution.
